# Glucose homeostasis is impaired in mice deficient for the neuropeptide 26RFa (QRFP)

**DOI:** 10.1101/816728

**Authors:** Mouna El Mehdi, Saloua Takhlidjt, Fayrouz Khiar, Gaëtan Prévost, Jean-Luc do Rego, Jean-Claude do Rego, Alexandre Bénani, Emmanuelle Nedelec, David Godefroy, Arnaud Arabo, Benjamin Lefranc, Jérôme Leprince, Youssef Anouar, Nicolas Chartrel, Marie Picot

## Abstract

**Introduction:** 26RFa (QRFP) is a biologically active peptide that has been found to control feeding behaviour by stimulating food intake, and to regulate glucose homeostasis by acting as an incretin. The aim of the present study was thus to investigate the impact of 26RFa gene knockout on the regulation of energy and glucose metabolism.

**Research design and methods:** 26RFa mutant mice were generated by homologous recombination, in which the entire coding region of prepro-26RFa was replaced by the iCre sequence. Energy and glucose metabolism was evaluated through measurement of complementary parameters. Morphological and physiological alterations of the pancreatic islets were also investigated.

**Results:** Our data do not reveal significant alteration of energy metabolism in the 26RFa-deficient mice except the occurrence of an increased basal metabolic rate. By contrast, 26RFa mutant mice exhibit an altered glycemic phenotype with an increased hyperglycemia after a glucose challenge associated with an impaired insulin production, and an elevated hepatic glucose production. 2D and 3D immunohistochemical experiments indicate that the insulin content of pancreatic β cells is much lower in the 26RFa^-/-^ mice as compared to the wild-type littermates.

**Conclusion:** Disruption of the 26RFa gene induces substantial alteration in the regulation of glucose homeostasis with, in particular, a deficit in insulin production by the pancreatic islets. These findings further support the notion that 26RFa is an important regulator of glucose homeostasis.

**Significance of this study:** *What is already known about this subject?:* 26RFa is a biologically active peptide produced in abundance in the gut and the pancreas. 26RFa has been found to regulate glucose homeostasis by acting as an incretin and by increasing insulin sensitivity.

*What are the new findings?:* Disruption of the 26RFa gene induces substantial alteration in the regulation of glucose homeostasis with, in particular, a deficit in insulin production by the pancreatic islets, assessing therefore the notion that 26RFa is an important regulator of glucose homeostasis.

*How might these results change the focus of research or clinical practice?:* Identification of a novel actor in the regulation of glucose homeostasis is crucial to better understand the general control of glucose metabolism in physiological and pathophysiological conditions, and opens new fields of investigation to develop innovative drugs to treat diabetes mellitus.

## Introduction

Obesity and diabetes are considered as epidemic worldwide issues. Indeed, in 2016, 650 million adults were classified as clinically obese (1), and 300 million people are affected by type 2 diabetes (T2DM) worldwide (http://www.idf.org/). Obesity and T2DM are two strongly associated diseases as 80% of people developing T2DM are obese, with insulin resistance as a common feature of both obesity and diabetes, raising therefore the hypothesis that obesity and diabetes may arise from a common functional defect.

Interestingly, accumulating data obtained during the last decade revealed that a number of neuropeptides well known to control feeding behaviour such as NPY, orexins, ghrelin, CRF or apelin, may also regulate glucose homeostasis (2-6). It was also found that the neuropeptidergic systems controlling feeding behaviour and glucose homeostasis in the hypothalamus partially overlap (7,8). From these observations suggesting that neuropeptides may link energy and glucose homeostasis, emerged a new concept proposing that the pathogenesis of obesity and diabetes may originate from defects of the neuropeptidergic systems controlling both energy and glucose homeostasis (9).

In this context, the neuropeptidergic system, the 26RFa/GPR103 system, is of particular interest. 26RFa (also referred to as QRFP) is a hypothalamic neuropeptide discovered concurrently by us and others (10-12). 26RFa has been characterized in all vertebrate phyla including human (13, 14), and identified as the cognate ligand of the human orphan G protein-coupled receptor, GPR103 (11-13,15, 16). Neuroanatomical observations revealed that 26RFa- and GPR103-expressing neurons are primarily localized in hypothalamic nuclei involved in the control of feeding behaviour (10,11,15,17,18). Indeed, i.c.v. administration of 26RFa stimulates food intake (10,15,19,20), and the neuropeptide exerts its orexigenic activity by modulating the NPY/POMC system in the arcuate nucleus (Arc) (20). 26RFa also stimulates food intake in birds (21) and fish (22), indicating that 26RFa plays a crucial role in the central regulation of body weight and energy homeostasis in all vertebrates. Interestingly, a more sustained orexigenic activity of 26RFa was reported in obese rodents (19,23). In addition, expression of prepro26RFa mRNA is up-regulated in the hypothalamus of genetically obese *ob/ob* and *db/db* mice (15), in rodents submitted to a high fat diet (19,23), and plasma levels of the neuropeptide are increased in obese patients (24,25). Finally, 26RFa was found to trigger lipid uptake and to inhibit lipolysis in obese individuals (26). Altogether, these findings support the notion that 26RFa could play a role in the development and maintenance of the obese status (14).

More recently, the implication of the 26RFa/GPR103 neuropeptidergic system in the control of glucose homeostasis was reported. We, and others, found that 26RFa and GPR103 are strongly expressed by β cells of the pancreatic islets (24,25,27), and that the neuropeptide prevents cell death and apoptosis of β cells (27). We also showed that 26RFa is abundantly expressed all along the gut and that i.p. administration of the neuropeptide attenuates glucose-induced hyperglycemia by increasing plasma insulin via a direct insulinotropic effect on the pancreatic β cells, and by increasing insulin sensitivity (24,25). Finally, we reported that an oral glucose challenge induces a massive secretion of 26RFa by the gut into the blood, strongly suggesting that this neuropeptide regulates glycemia by acting as an incretin (24). This incretin effect of 26RFa has been very recently confirmed by the observation that administration of a GPR103 antagonist reduces the global glucose-induced incretin effect, and also decreases insulin sensitivity (28).

Together, these findings support the idea that the 26RFa/GPR103 peptidergic system plays an important role in the regulation of energy and glucose homeostasis. Consequently, the objective of the present study was to investigate the impact of altered endogenous 26RFa production on the regulation of energy and glucose metabolism using a model of mice invalidated for the 26RFa gene.

## Research Design and Methods

### Animals

26RFa^-/-^, 26RFa^+/-^ and 26RFa^+/+^ male C57Bl/6 mice, weighing 22–25 g, were housed with free access to standard diet (U.A.R., Villemoisson-sur-Orge, France) and tap water. They were kept in a ventilated room at a temperature of 22±1°C under a 12-h light/12-h dark cycle (light on between 7 h and 19 h). All the experiments were carried out between 09.00 h and 18.00 h in testing rooms adjacent to the animal rooms. Mice were housed at three to five per cage. Unless otherwise stated, all tests were conducted with naïve cohorts of mice that were habituated to physiological and behaviour protocols before the beginning of experiments.

All experimental procedures were approved by the Normandy Regional Ethics Committee (Authorization: APAFIS#11752-2017100916177319) and were carried out in accordance with the European Committee Council Directive of November 24, 1986 (86/609/EEC).

### Metabolic phenotype analysis

#### Combined indirect calorimetry

A 16-cage combined indirect calorimetry system (PhenoMaster, TSE Systems GmbH, Bad Homburg, Germany) was used to assess continuous monitoring of the energy expenditure, locomotor activity, respiratory quotient as well as food and water intake. Mice were individually housed and acclimated to the air-tight cages for 5 days before experimental measurements. Subsequently, volumes of oxygen consumption and volumes of CO_2_ production were measured every minute for a total of six light and six dark phases (144 h) to determine the respiratory quotient (RQ = VCO_2_/VO_2_) and energy expenditure (EE = VO_2_ × (3.815+(1.232 × (VCO_2_/VO_2_))) × 4.1868).

16 Home-cage locomotor activity (horizontal and vertical) was determined by a multidimensional infrared light beam system. Stationary locomotor activity was defined as consecutive infrared light beam breaks of one single light beam and ambulatory movement as consecutive breaks of two different light beams.

Scales integrated into the sealed cage environment continuously measured cumulative food intake and water intake. The mice were kept at a constant temperature of 23°C for 6 days.

#### Body composition

Whole body composition was assessed on vigil animals, before and after metabolic parameters measurement, using MiniSpec LF110 (Brucker, Wissembourg, France), a fast nuclear magnetic resonance method.

### Blood glucose and insulin measurements in mice

For measurements of basal glycemia and insulinemia, mice were fasted 6 h before the test with free access to water. For oral or i.p. glucose tolerance test, mice were fasted for 16 h with free access to water and then treated i.p. or by gavage with glucose (2 g/kg) and 26RFa (500 μg/kg) for reversion experiments. For insulin tolerance test, mice were fasted for 6 h before the test with free access to water, and then injected i.p. with 0.75 units/kg body weight of human insulin (Eli Lilly, Neuilly-sur-Seine, France). For pyruvate tolerance test, mice were fasted for 16 h before the test with free access to water and then injected i.p. with sodium pyruvate (2 g/kg; Sigma Aldrich) and 26RFa (500 μg/kg) for reversion experiments. Plasma glucose concentrations were measured from tail vein samplings at various times using an AccuChek Performa glucometer (Roche Diagnostic, Saint-Egreve, France). Plasma insulin concentrations were determined using an ultrasensitive mouse insulin AlphaLisa detection kit (cat number AL204C) from Perkin Elmer.

### Quantitative PCR

Total RNA from livers of mice was isolated as previously described (28). Relative expression of the glucose 6 phosphatase (G6PC), glucokinase (GCK) and phosphoenolpyruvate carboxykinase 1 (PCK1) genes was quantified by real-time PCR with appropriate primers (Table 1). β-actin was used as internal control for normalization. PCR was carried out using Gene Expression Master Mix 2X assay (Applied Biosystems, Courtaboeuf, France) in an ABI Prism 7900 HT Fast Real-time PCR System (Applied Biosystems). The purity of the PCR products was assessed by dissociation curves. The amount of target cDNA was calculated by the comparative threshold (Ct) method and expressed by means of the 2-ΔΔCt method.

**Table 1:**
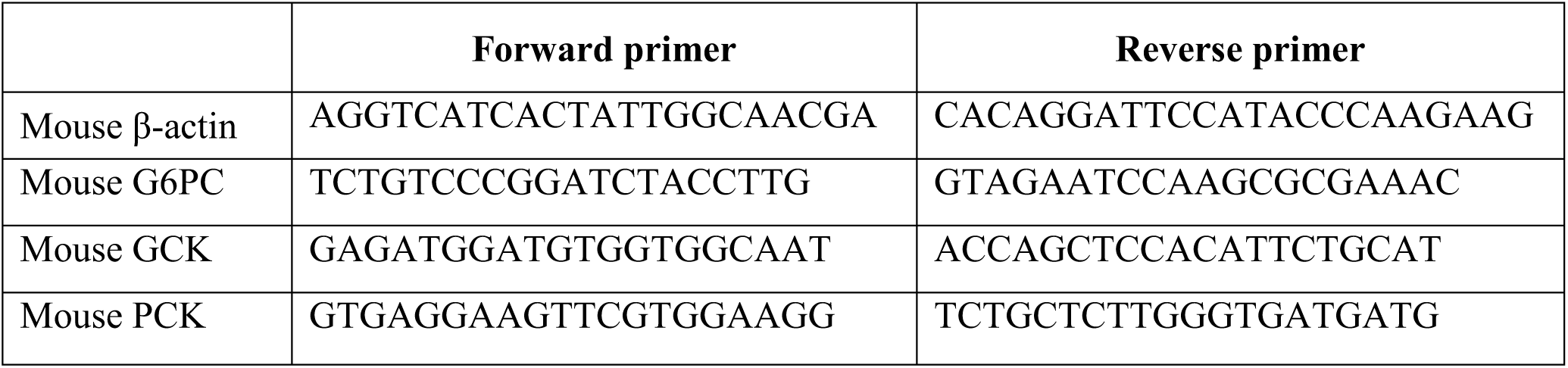
Sequence of the primers used for the Q-PCR experiments

### Morphological analysis of the pancreas

Deparaffinized sections (15-µm thick) of 26RFa^-/-^, 26RFa^+/-^ and 26RFa^+/+^ mice pancreas and duodenum were used for immunohistochemistry. For the observation of the 26RFa immunolabelling, tissue sections were incubated for 1 h at room temperature with rabbit polyclonal antibodies against 26RFa (25) diluted 1:400. The sections were incubated with a streptavidin-biotin-peroxydase complex (Dako Corporation, Carpinteria, CA), and the enzymatic activity was revealed with diaminobenzidine. Some slices were then counterstained with hematoxylin. Observations were made under a Nikon E 600 light microscope. For the study of the morphological architecture, pancreas slices were stained with hematoxylin/eosin and examined under a Leica Z6 APO macroscope and a Nikon E 600 light microscope.

For the double labelling experiments, the following primary antibodies were used: guinea pig polyclonal anti-insulin (1:50, Gene Tex, Irvine, CA) and mouse monoclonal anti-glucagon (1:1000, Sigma Aldrich). Alexa-conjugated antibodies (Invitrogen Life Technologies) including goat anti-guinea pig-488 (1:300) and donkey anti-mouse-594 (1:300) were used as secondary antibodies. Sections were counterstained with 1 μg/mL 4′,6-diamino-2-phenylindole (Sigma-Aldrich) in PBS for 90 seconds. Tissue sections were examined with a Nikon Eclipse E600. Quantitative, qualitative and morphological analysis of the pancreas sections were performed using the Image J software (NIH, Washington, DC).

### 3D pancreas analysis

Treatment of whole pancreas of 26RFa^-/-^and 26RFa^+/+^mice for immunohistochemistry *in toto* was performed according to a protocol previously published (29). Briefly, the pancreas were fixed by perfusion of PFA 4%. After dehydratation in successive baths of methanol solution (20%, 40%, 60%, 80% in PBS 1X and 2X100%, 1h each), bleaching in H_2_O_2_ and rehydratation with successively methanol solution (100%, 80%, 60%, 40%, 20% and PBS1X, 1 h each), a step of permeabilization was performed by incubating the tissues in a permeabilization solution containing 0.2% Triton X-100, glycine 0.3 M, 20% DMSO for 4 days with rotation at room temperature. Then, an antigen blocking step was performed by incubating the tissues in PBS containing 0.2% Triton X-100, 10% DMSO and 6% donkey serum for 2 days at 37°C. Pancreas were incubated in a PBS solution with 0,2% tween 20, 0,1% heparin (10mg/ml), 5% DMSO, 3% donkey serum with mouse monoclonal insulin antibodies (Sigma-Aldrich) at a dilution of 1:200 for 11 days, and then for 6 days with a donkey anti-mouse IgG (FP-SC4110-E, Interchim) diluted 1:300, used as secondary antibody. Pancreas were cleared with final steps of iDISCO+ protocol and processed for imaging. For this, the immunolabeled pancreas were visualized in three dimensions using an Ultramicroscope II (Light Sheet Microscope; LaVision BioTec, Bielefeld, Germany) equipped with a Neo sCMOS camera (Andor Technology, Belfast, UK). 3D reconstructions were made with the Imaris Software version 8.4 (Bitplane, Zurich, Switzerland).

### Statistical analysis

Statistical analysis was performed with GraphPad Prism (6^th^ version). A student *t*-test or an ANOVA one way were used for comparison between the groups. An ANOVA two ways was used for repeated measures for comparisons between the groups. A post-hoc comparison using Tukey HSD was applied according to the ANOVA two ways results. Statistical significance was set up at p < 0.05.

## Results

### Strategy and characterization of mouse 26RFa gene disruption

26RFa^-/-^ (iCre knock-in) mice were obtained from Prof T. Sakurai (International Institute for Integrative Sleep Medecine, Tsukuba, Ibaraki, Japan). The mutant mice were generated by homologous recombination in embryonic stem cells of 129SvJ strain and implanted in C57 blastocysts using standard procedures. The targeting vector was constructed by replacing the entire coding region of prepro-26RFa sequence in the exon 2 of the 26RFa gene with iCre sequence and pgk-Neo cassette (Suppl data 1A).

Genotypes were determined by PCR of DNA mouse tail biopsy. PCR primers used were 5’-CAGTCAGCAGCTATCCCTCC-3’ (from 115 to 96 base of the 26RFa gene from transcription initiation site) and 5’-ACCGTCTTGCCTCCCTAGACG-3’ (from 225 to 246 base), and 5’-GAGGGACTACCTCCTGTACC-3’ and 5’-TGCCCAGAGTCATCCTTGGC-3’ (Corresponding to the iCre sequence). We detected a 361-pb product from wildtype allele corresponding to the 26RFa coding sequence, and a 650-pb product from the targeted allele corresponding to the inserted iCre sequence (Suppl data 1B). Chimeric mice were crossed with C57Bl/6J males (Janvier laboratory, Le Genest-Saint-Isle, France). Initially, F1 hybrids from heterozygous x heterozygous mating were generated. 26RFa^-/-^ mice and wild type mice littermates were basically obtained by heterozygous x heterozygous mating.

Immunohistochemical experiments performed on pancreas and duodenal sections revealed the presence of an intense 26RFa-immunolabelling in the pancreatic islets and the duodenal enterocytes of the wild type mice whereas similar sections from 26RFa-deficient mice were totally devoid of staining (Fig. 1A).

**Figure 1.**
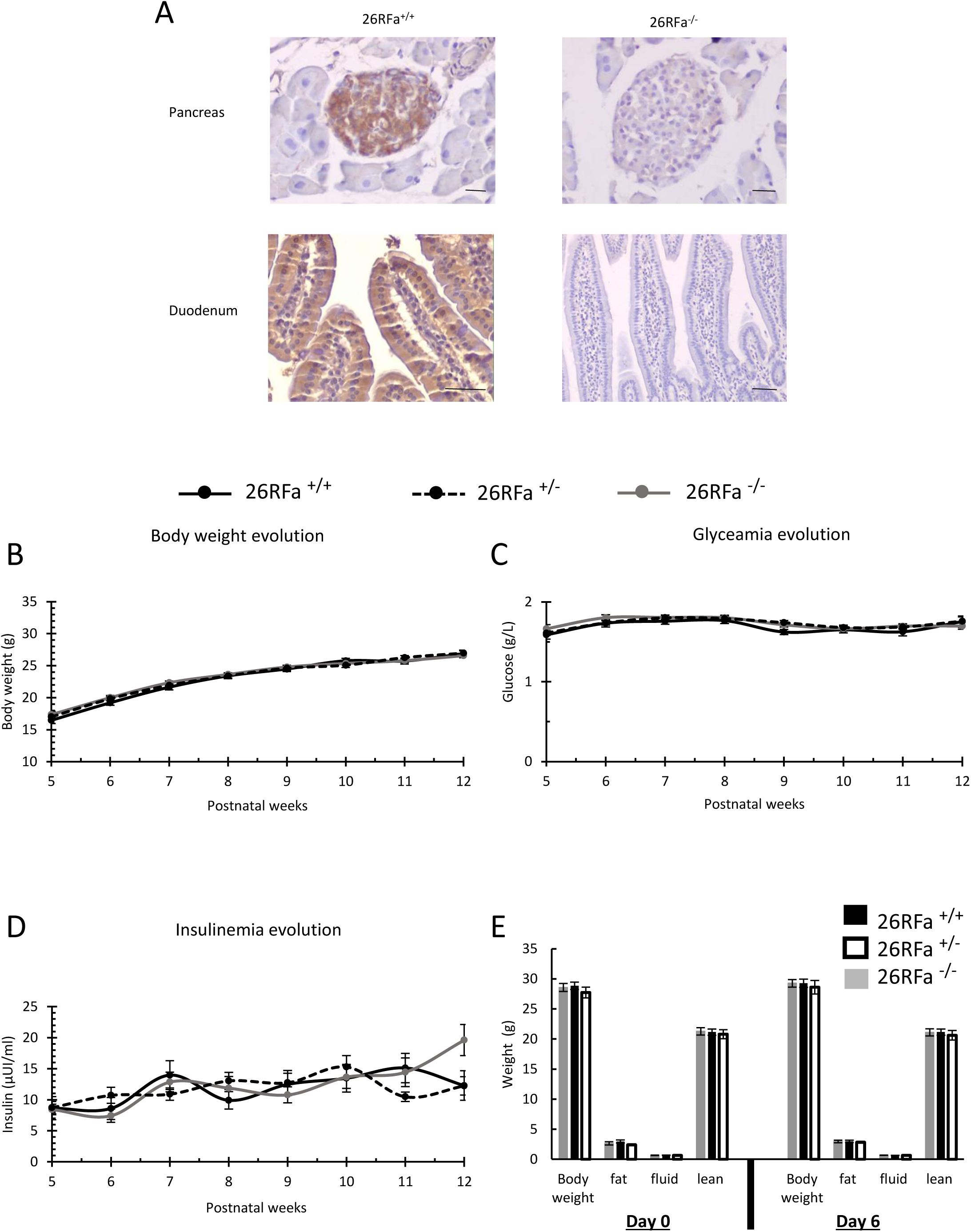
Characterization and body composition of 26RFa-deficient mice. **A:** Immunohistochemical photomicrographs of pancreas and duodenal sections showing complete depletion of the 26RFa immunostaining in the tissues of the 26RFa^-/-^ mice. Scale bars: 50 µm. **B-D:** Evolution of body weight, glycemia and insulinemia from the 5^th^ to the 12^th^ postnatal week (n=20-58). **E:** Evaluation of body mass composition in 26RFa^-/-^, 26RFa^+/-^ and 26RFa^+/+^ mice during a 6-day experimental protocol in which animals were fed ad libitum for 3 days, then were food-restricted for one day and refed for two days (n=11 per group). Data represent means ± SEM.

### Energy metabolism phenotype of 26RFa-deficient mice

Measurement of body weight, plasma glucose and insulin levels from post-natal week 5 to week 12 revealed no significant difference between the wild type, heterozygous and 26RFa-KO mice (Fig. 1B-D). Two-month old mice of the 3 groups also followed a 6-day protocol in which the animals were fed ad libitum for 3 days, then were food-restricted for one day and refed for two days. The body composition analysis showed that 26RFa^-/-^, 26RFa^+/-^ and 26RFa^+/+^mice had similar body weight, fat mass, lean mass and fluid mass, either at the beginning of the cession or at the end of the protocol (Fig. 1E).

Measurement of food intake all along the protocol did not reveal any significant difference in the feeding behaviour of the 3 groups of mice although the 26RFa-KO and the heterozygous mice ate a little bit more than the wild type controls (Fig. 2A). Evaluation of water intake indicated the 26RFa^-/-^ mice drink significantly more than the wild type and the heterozygous mice during the test (Fig. 2B). Measurement of energy expenditure did not reveal any significant variation between the 3 groups of mice although the 26RFa-deficient mice exhibited a tendency to increased energy expenditure as compared to the wild type and heterozygous mice (Fig. 2C). Metabolic rate measured as O_2_, CO_2_ and respiratory quotient was significantly higher in 26RFa-deficient mice as compared to wild type and heterozygous mice (Fig. 2D-F). The locomotor activity (horizontal and vertical) was also measured but did not reveal any significant difference between the 26RFa^-/-^, 26RFa^+/-^ and 26RFa^+/+^ mice (Fig. 2G, H).

**Figure 2.**
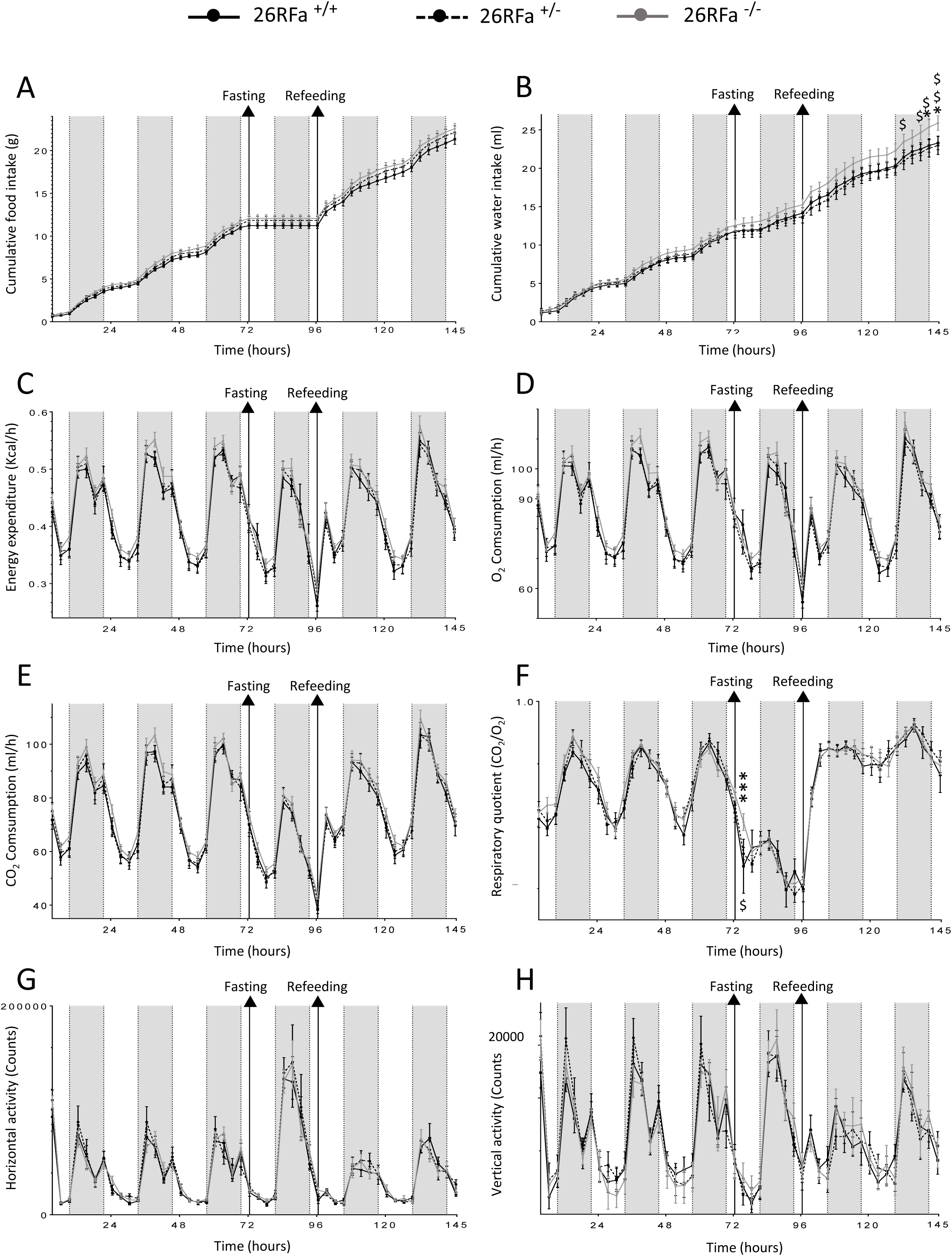
Energy metabolism phenotype of 26RFa-deficient mice. **A-F:** Evaluation of various metabolic parameters including feeding (A) and drinking (B) consumption, energy expenditure (C), O_2_ consumption (D), CO_2_ consumption (E) and respiratory quotient (F) in 26Rfa^-/-^, 26RFa^+/-^ and 26RFa^+/+^ mice during a 6-day experimental protocol in which animals were fed ad libitum for 3 days, then were food-restricted for one day and refed for two days (n=11 per group).**G, H:** Evaluation of locomotor activity in 26RFa^-/-^, 26RFa^+/-^ and 26RFa^+/+^ mice during the same experimental protocol as in (A-F) (n=11 per group). Data represent means ± SEM. *, p<0.05; ***, p<0.001 26RFa^-/-^ *vs* 26RFa^+/+^ mice. $, p<0.05; $$, p<0.01 26RFa^-/-^ *vs* 26RFa^+/-^ mice.

### Glycemic phenotype of 26RFa-deficient mice

The “glycemic” phenotype of the 26RFa-deficient and the heterozygous mice was investigated using complementary *in vivo* tests. Basal plasma glucose levels after a 6h fasting were comparable in the 26RFa^-/-^, 26RFa^+/-^ and 26RFa^+/+^ mice (Fig. 3A). By contrast, basal plasma insulin levels were significantly lower in the 26RFa-deficient mice as compared to the wild type and the heterozygous mice (Fig. 3B). An oral glucose tolerance test (OGTT) indicated that the hyperglycemic and hyperinsulinemic peaks induced by the glucose load were not affected by the lack of 26RFa (Fig. 3C, D). Conversely, the intraperitoneal glucose tolerance test (IPGTT) revealed a more sustained hyperglycemic peak in the 26RFa^-/-^ mice that was associated with a lower rise of plasma insulin levels (Fig. 3E, F). The heterozygous mice exhibited a glycemic and insulinemic profile during the IPGTT between those of the wild type and KO animals (Fig. 3E, F). I.p. administration of 26RFa totally reversed the alterations in plasma glucose and insulin observed during the IPGTT in the 26RFa-deficient mice (Fig. 3G, H).

**Figure 3.**
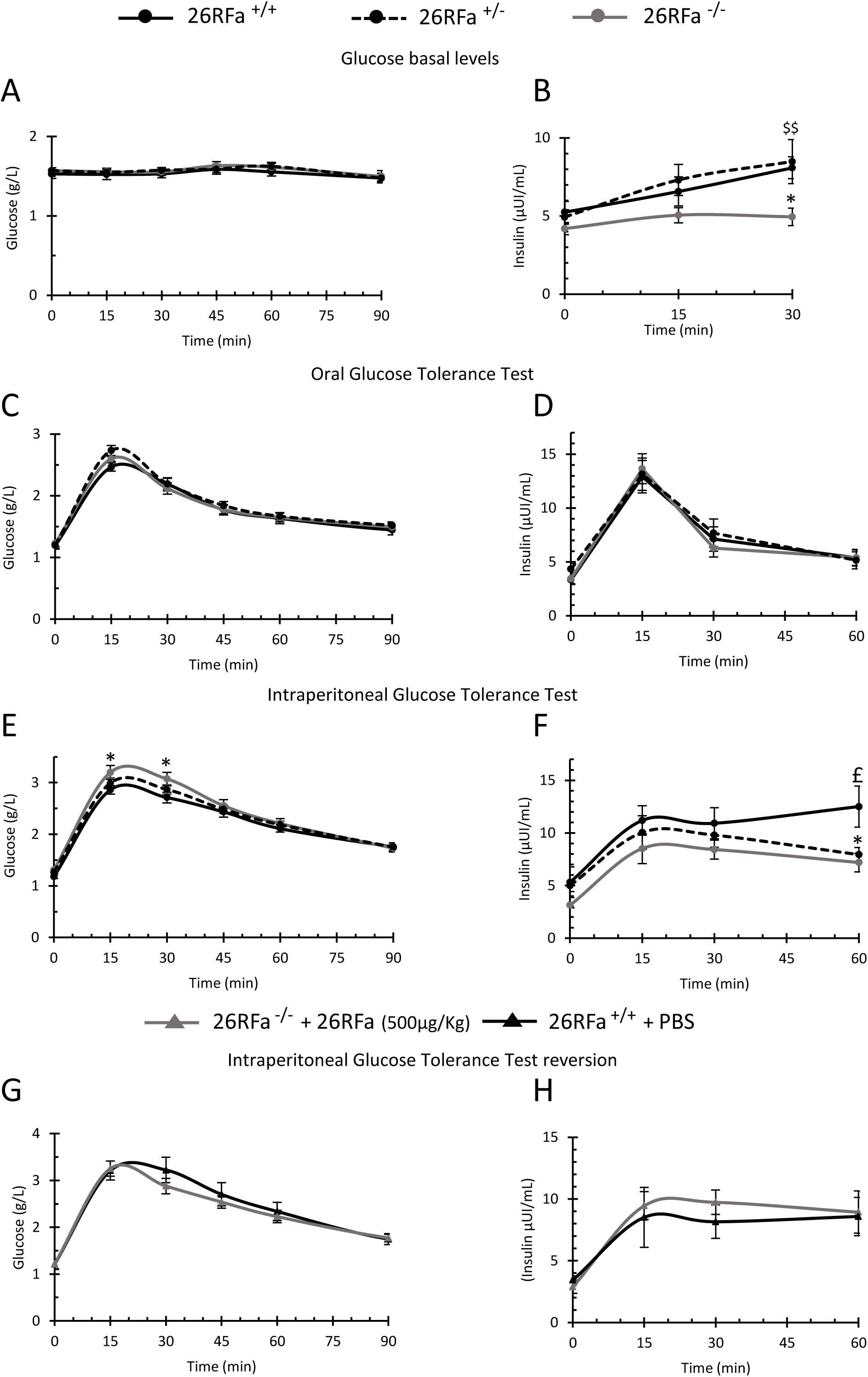
Glycemic phenotype of 26RFa-deficient mice 1. **A, B:** Evaluation of plasma glucose and insulin levels in 26RFa^-/-^, 26RFa^+/-^ and 26RFa^+/+^ mice in basal condition (n=10-18 per group). **C, D:** Evaluation of plasma glucose and insulin levels in 26RFa^-/-^, 26RFa^+/-^ and 26RFa^+/+^ mice during an oral glucose tolerance test (n=12-17 per group). **E, F:** Evaluation of plasma glucose and insulin levels in 26RFa^-/-^, 26RFa^+/-^ and 26RFa^+/+^ mice during an intraperitoneal glucose tolerance test (n=12-17 per group). **G, H:** Evaluation of plasma glucose and insulin levels in 26RFa^-/-^ mice that received an i.p. dose of 26RFa (500 µg/kg) during an intraperitoneal glucose tolerance test (n=8 per group). Data represent means ± SEM of 4 independent experiments. *, p<0.05 26RFa^-/-^ *vs* 26RFa^+/+^ mice. $$, p<0.01 26RFa^-/-^ *vs* 26RFa^+/-^ mice.

The impact of the 26RFa gene disruption on insulin sensitivity and hepatic glucose production was also examined. An insulin tolerance test (ITT) revealed that insulin sensitivity was not altered in the 26RFa^-/-^ mice and the 26RFa^+/-^ mice as compared to the wild type animals (Fig. 4A). By contrast, a pyruvate tolerance test (PTT) showed that hepatic glucose production was significantly increased in the 26RFa-KO mice in comparison to the wild type animals (Fig. 4B). I.p. administration of 26RFa in the 26RFa^-/-^ mice reversed the hyperglycemia observed during the PTT (Fig. 4C). In addition, expression of liver enzymes playing a key role in gluconeogenesis and glucogenolysis was determined after a 16-h fasting that promotes glucose hepatic production, and was compared to fed condition. As expected, in fasting condition, wild type animals showed a drastic decrease of glucokinase (GCK) that promotes glycogen storage and an upregulation of glucose 6 phosphatase (G6PC) and phosphoenolpyruvate carboxykinase 1 (PCK1) that trigger gluconeogenesis (Fig. 4D). The 26RFa-deficient mice exhibited a different expression profile in fasting condition with a slight decreased expression of GCK associated with a robust increased expression of G6PC and PCK1 (Fig. 4E).

**Figure 4.**
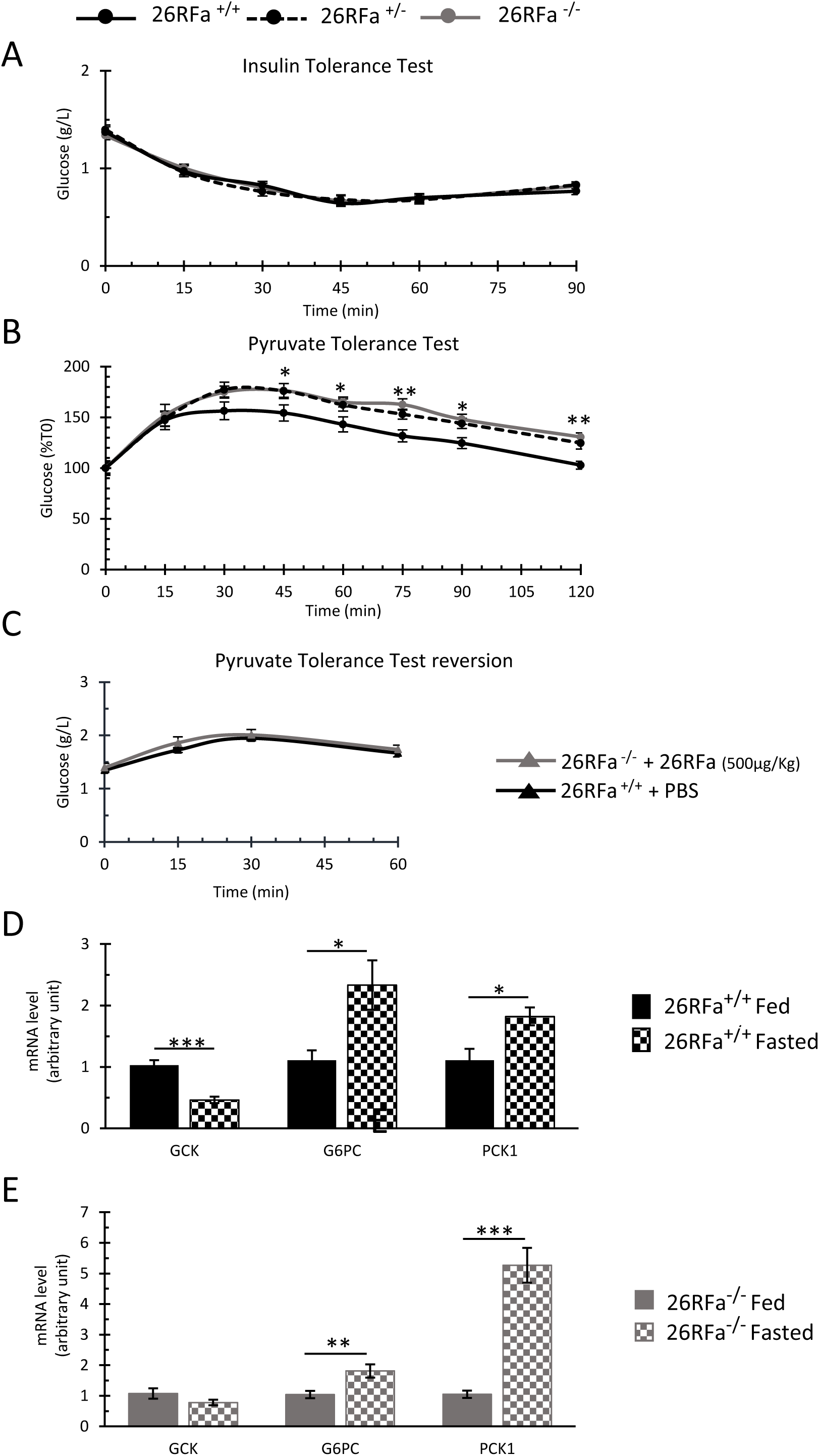
Glycemic phenotype of 26RFa-deficient mice 2. **A:** Evaluation of plasma glucose levels in 26RFa^-/-^, 26RFa^+/-^ and 26RFa^+/+^ mice during an insulin tolerance test (n=13-18 per group). **B:** Evaluation of plasma glucose levels in 26RFa^-/-^, 26RFa^+/-^ and 26RFa^+/+^ mice during a pyruvate tolerance test (n=10-14 per group). **C:** Evaluation of plasma glucose levels in 26RFa^-/-^ mice that received an i.p. dose of 26RFa (500 µg/kg) during a pyruvate tolerance test (n=8 per group). **D, E:** expression of the liver enzymes glucokinase (GCK), glucose 6 phosphatase (G6PC) and phosphoenolpyruvate carboxykinase 1 (PCK1) in fasted or fed conditions of 26RFa^+/+^(D) and 26RFa^-/-^ mice (E) (n=8 per group). Data represent means ± SEM of 3 independent experiments. *, p<0.05; **, p<0.01; ***, p<0.001 26RFa^-/-^ *vs* 26RFa^+/+^ mice.

### Pancreatic phenotype of 26RFa-deficient mice

Comparison of freshly dissected pancreas indicated that the tissues of the 26RFa-KO mice were bigger with more adipose tissues than the wild type and heterozygous mice although their weights were similar in the 3 groups (Fig. 5A, B). We also observed that the number of pancreatic islets per pancreas tended to be higher in the 26RFa^+/-^ and 26RFa^-/-^ mice, although statistically not significant (Fig. 5C). In addition, the areas of the pancreatic islets were significantly higher in the 26RFa-deficient mice as compared to the wild type mice (Fig. 5D), as illustrated by the photomicrographs shown in figure 5E. The quantitative analysis also revealed that the total number of β cells per pancreas was significantly higher in the 26RFa^-/-^ mice *vs* 26RFa^+/+^ mice (Fig. 5F). Conversely, the number of α cells per islet was significantly lower in the 26RFa^+/-^ and 26RFa^-/-^ mice *vs* 26RFa^+/+^ mice (Fig. 5G). Consequently, the ratio α cells/β cells per islet was significantly lower in the heterozygous and KO mice as compared to the wild type animals (Fig. 5H). Triple labelling experiments revealed that the intensity of the insulin and glucagon immunostaining was much lower in the 26RFa-deficient mice *vs* wild type animals, whereas, in the heterozygous mice the intensity of the immunostaining of the two hormones was between the 26RFa^-/-^ and the 26RFa^+/+^ mice (Fig. 5I). The iDISCO approach confirmed that the intensity of the insulin immunostaining was much higher in the wild type animals (video S1) in comparison to the 26RFa-KO mice (video S2).

**Figure 5.**
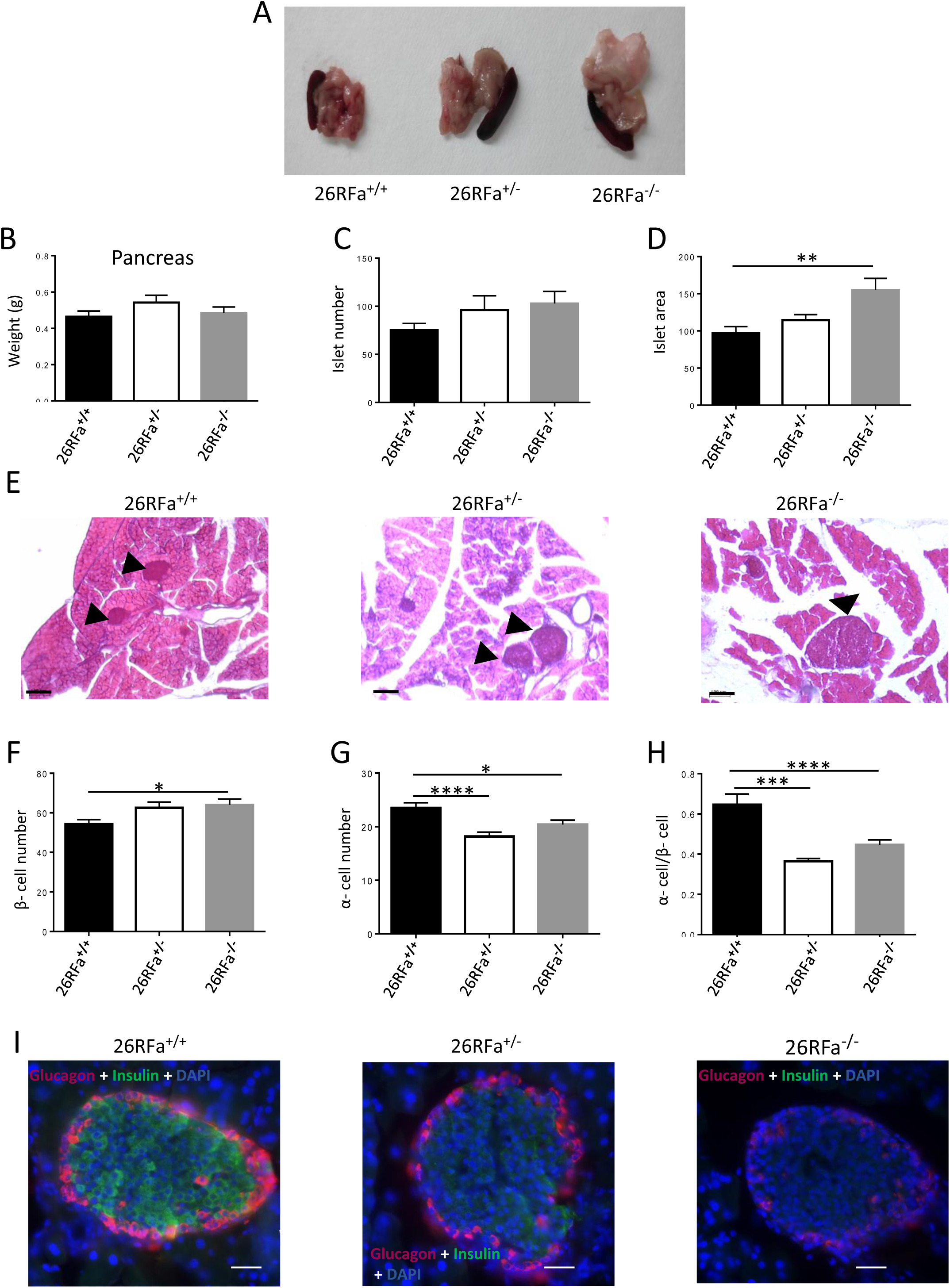
Pancreatic phenotype of 26RFa-deficient mice. **A, B:** Representative photomicrographs of freshly dissected pancreas of 26RFa^-/-^, 26RFa^+/-^ and 26RFa^+/+^ mice with their respective weights (n=8 per group). **C, D:** Quantitative analysis of the number of pancreatic islets and their areas in 26RFa^-/-^, 26RFa^+/-^ and 26RFa^+/+^ mice (n=5 per group). Data represent means ± SEM. **, p<0.01. **E:** Representative photomicrographs showing the difference in the size of the pancreatic islets 26RFa^-/-^, 26RFa^+/-^ and 26RFa^+/+^mice (arrowheads). **F-H:** Quantitative analysis of the number of β and α cells per pancreatic islet and their ratio in 26RFa^-/-^, 26RFa^+/-^ and 26RFa^+/+^ mice (n=5 per group). **I:** Representative photomicrographs of pancreatic islets of 26RFa^-/-^, 26RFa^+/-^ and 26RFa^+/+^ mice labelled with an insulin antibody (green), a glucagon antibody (red) and DAPI (blue), showing that the intensity of the insulin and glucagon immunostaining is much lower in the 26RFa^-/-^ mice than in the 26RFa^+/+^ mice. Data represent means ± SEM. *, p<0.05; ***, p<0.001. Scale bars: 100 µm.

## Discussion

Accumulated data obtained during the last decade have promoted the evidence that the neuropeptide 26RFa plays a key role in the control of feeding behaviour (10, 13, 14) and the regulation of glucose homeostasis (24, 25, 27, 28). Supporting this notion, it has been recently shown that acute administration of a GPR103 (the 26RFa receptor) antagonist decreases food intake (30) and reduces the global glucose-induced incretin effect as well as the insulin sensitivity (28). However, chronic deficiency of 26RFa signalling on energy and glucose homeostasis remains to be elucidated. In the present study, we took advantage of a newly generated mouse line deficient for the 26RFa gene to decipher the phenotype of the 26RFa^-/-^mice with regard to energy and glucose metabolism.

We first investigated the impact of chronic 26RFa depletion on various parameters of energy metabolism. Our data reveal that 26RFa deficiency does not alter body weight gain from postnatal week 5 to week 12. We also show that at 2 months, the 26RFa^-/-^, 26RFa^+/-^ and 26RFa^+/+^ mice exhibit a similar body composition in terms of body weight, fat and lean mass. However, the 26RFa-KO mice show a basal energy expenditure slightly higher than the heterozygous and their wild type littermates. In addition, the 26RFa-deficient mice tend to eat and drink more than the wild type mice and exhibit a more elevated respiratory quotient although their locomotor activity is not altered. Collectively, these observations suggest that 26RFa-deficient mice have a basal metabolic rate slightly higher than the wild type animals. The observation that deletion of the 26RFa gene does not impair daily feeding behaviour and body weight is not surprising as disruption of other major orexigenic peptides such as NPY or ghrelin does not impact feeding behaviour (31, 32). Indeed, it is accepted that the congenital lack of one regulatory peptide may be compensated by others as the control of feeding behaviour is multifactorial. However, our data are in disagreement with a previous study reporting that disruption of the 26RFa (QRFP) gene results in a lower body weight due to a hypophagic behaviour under both normal and high fat fed condition (33). We do not have any obvious explanation for this discrepancy between the two studies except that the strains of 26RFa^-/-^mice in the two studies are different (33, present study).

In the second part of our study, we have investigated the “glycemic” phenotype of the 26RFa-deficient mice. We first show that depletion of the 26RFa gene has no impact on the evolution of glycemia and insulinemia measured in fed condition. Basal plasma glucose levels are not altered in fasting condition in the 26RFa-KO mice too. However, fasted insulinemia is significantly decreased in the 26RFa^-/-^ mice as compared to the 26RFa^+/-^and 26RFa^+/+^ mice. In addition, we show that, during a glucose tolerance test, the hyperglycemic peak is more sustained in the mice deficient for 26RFa and this is associated with a lower rise in plasma insulin, these effects being reversed by administration of exogenous 26RFa. By contrast, our data indicate that insulin sensitivity is not affected by the absence of endogenous 26RFa. Altogether, these findings reveal that depletion of 26RFa induces an alteration of glucose homeostasis that is due to a defect in insulin secretion but not in insulin sensitivity. Consistent with this finding, we have previously shown that 26RFa stimulates insulin secretion by the pancreatic β cells (24) and that administration of a 26RFa receptor antagonist alters the anti-hyperglycemic response of the organism to a glucose challenge (28). Collectively, these data confirm that 26RFa is an important regulator of glucose homeostasis.

Our data also reveal that depletion of 26RFa induces a dysfunction in the regulation of glucose hepatic production. Indeed, the 26RFa-deficient mice exhibit an increase of their glucose hepatic production that is associated with an up-regulation of G6PC and PCK1, two key liver enzymes that trigger gluconeogenesis. We have previously reported that 26RFa exerts a crucial anti-hyperglycemic effect that is due to its incretin activity and its insulin-sensitiviting effect (24). Our present finding suggests that the inhibitory activity of 26RFa on hepatic glucose production also participates to the global anti-hyperglycemic effect of the peptide. However, we have also previously shown that the 26RFa receptor GPR103 is not expressed in the liver (24). This observation suggests that the inhibitory effect of 26RFa on glucose hepatic production is not a direct effect but rather an indirect effect, maybe mediated *via* its insulin secreting activity as insulin is well known to inhibit hepatic glucose production.

One major phenotype of the 26RFa-deficient mice is the low plasma insulin levels in fasting conditions or in response to a glucose challenge. This led us to examine whether this decreased insulin production was associated with alteration in the morphology and physiology of the pancreas. Surprisingly, our quantitative analysis revealed that the pancreas of the mutant mice are bigger with larger pancreatic islets than their wild type littermates with a higher number of β cells/islet, that rather suggests an increased capacity of the pancreatic islets to produce and secrete insulin in the 26RFa-deficient mice. However, 2D and 3D immunohistochemical labeling of the pancreatic islets with insulin antibodies shows that the intensity of insulin immunostaining in the β cells of the the 26RFa^-/-^ mice is much lower than that observed in 26RFa^+/+^ mice, suggesting that the capacity of the β cells to produce/secrete insulin is impaired in the mutant mice. We hypothesize that this altered insulin production may explain the lower plasma insulin levels observed in the 26RFa-deficient mice and that the increased number of pancreatic islets and β cells observed in the mutant mice may reflect a compensatory mechanism of the organism to counterbalance the low capacity of the β cells to produce insulin. We have previously shown in human (25) and rodents (24) that the β cells of the pancreatic islets highly express 26RFa. According to this latter observation, we think that a potential role for 26RFa in the synthesis/production of insulin within the β cells deserves further investigation.

Finally, our experiments indicate that the heterozygous mice have an intermediate “glycemic” phenotype between those of the mutant and the wild type animals.

In conclusion, the present study reveals that depletion of the 26RFa gene induces a substantial alteration in the regulation of glucose homeostasis with, in particular, a deficit in insulin production by β cells of the pancreatic islets. These original data confirm and confirm our previous studies (23, 24), supporting the idea that the neuropeptide 26RFa is a key regulator of glucose homeostasis and that dysfunction of the 26RFa/GPR103 peptidergic system may promote diabetes.

## Author Contributions

M.E.M., M.P. and N.C. contributed to the study design and interpretation, and wrote the manuscript. J.L.D.R, J.C.D.R, F.K., S.T., A.A., M.P. and M.E.M. performed the in vivo experiments on mice. S.T., F.K., D.G. and M.P. contributed to the immunohistochemical experiments and their quantitative analysis. F.K. and M.E.M. contributed to the PCR experiments. A. B. and E. N. performed the insulin assays and J. L. produced synthetic 26RFa. G.P. and Y. A. revised and approved the final version of the manuscript. N.C. is the guarantor of this work and, as such, had full access to all the data in the study and takes responsibility for the integrity of the data and the accuracy of the data analysis.

