## Supplemental Fig 1 and legends for "Glucose homeostasis is impaired in mice deficient for the neuropeptide 26RFa (QRFP)"

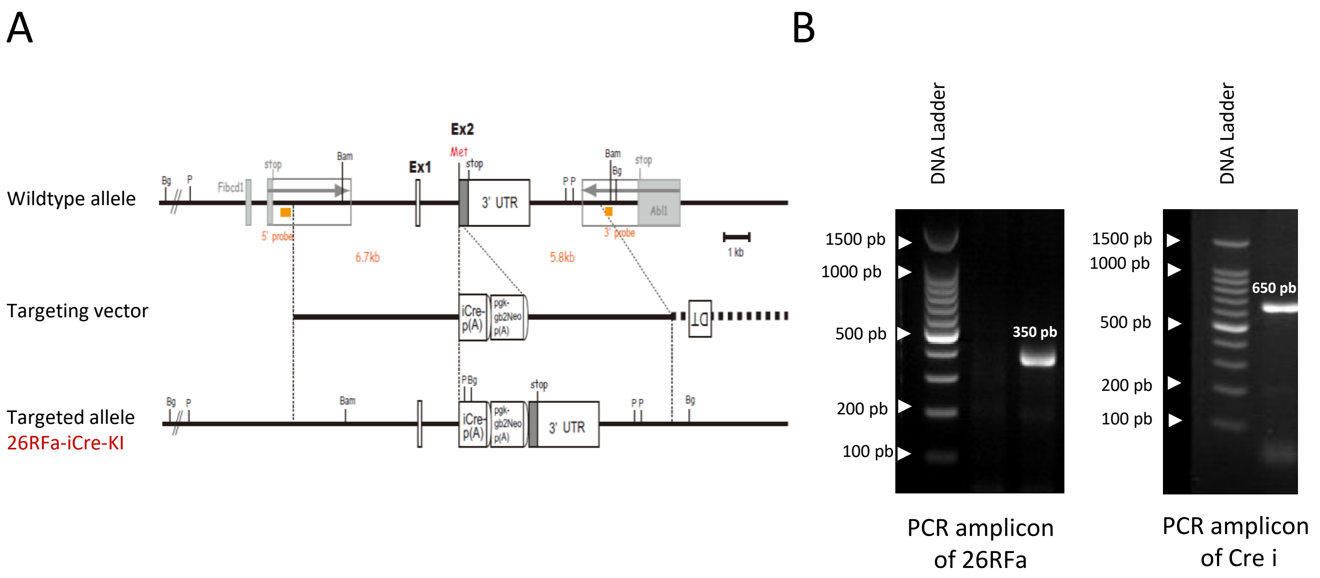


Supplementary data 1 :

### Legends to supplemental figures

**Supplemental data 1.** Strategy and characterization of mouse 26RFa gene disruption. **A:** Strategy for 26RFa disruption. Fibcd: fibrinogen C domain containing 1; Abl1: Proto-Oncogene 1, Non-Receptor Tyrosine Kinase; The PGK-gb2Neo template encodes the neomycin/kanamycin resistance gene which combines a prokaryotic promoter (gb2) for expression of kanamycin resistance and a eukaryotic promoter of phosphoglucokinase gene (PGK) for expression of neomycin resistance in mammalian cells. **B:** Genotyping by PCR of mouse tail DNA. In wild type mice, a 361-pb product corresponding to the 26RFa coding sequence was detected whereas, in 26RFa$^{-/-}$ mice, a 650-pb product corresponding to the inserted iCre sequence was found.

**Supplemental video S1 and S2.** Three-dimensional imaging of pancreas of 26RFa$^{+/+}$ (S1) and 26RFa$^{-/-}$ mice (S2) immunolabelled with an insulin antibody highlighting the important discrepancy in the intensity of the insulin immunostaining between the two genotypes. Original magnification, X4 (LaVision Ultramicroscope II Light Sheet Microscope).
